# Shotgun proteomic profiling of dormant, ‘non-culturable’ *Mycobacterium tuberculosis*

**DOI:** 10.1101/2021.08.06.455493

**Authors:** V. Nikitushkin, M. Shleeva, D. Loginov, F. Dycka, J. Sterba, A. Kaprelyants

## Abstract

Dormant cells of *Mycobacterium tuberculosis*, in addition to low metabolic activity and a high level of drug resistance, are characterized by ‘non-culturability’ – a specific reversible state of the inability of the cells to grow on solid media. The biochemical characterization of this physiological state of the pathogen is only superficial, pending clarification of the metabolic processes that may exist in such cells. In this study, applying LC-MS proteomic profiling, we report the analysis of proteins accumulated in dormant, ‘non-culturable’ *M. tuberculosis* cells in an *in vitro* model of self-acidification of mycobacteria in the post-stationary phase, simulating the *in vivo* persistence conditions. This approach revealed the accumulation of a significant number of proteins (1379) in cells after 4 months of storage in dormancy; among them, 468 proteins were significantly different from those in the actively growing cells and bore a positive fold change (FC). Differential analysis revealed the proteins of the pH-dependent regulatory system phoP and allowed the reconstruction of the reactions of central carbon/glycerol metabolism, as well as revealing the salvaged pathways of mycothiol and UMP biosynthesis, establishing the cohort of survival enzymes of dormancy. The annotated pathways mirror the adaptation of the mycobacterial metabolic machinery to life within lipid-rich macrophages, especially the involvement of the methyl citrate and glyoxylate pathways. Thus, the current *in vitro* model of *M. tuberculosis* self-acidification reflects the biochemical adaptation of these bacteria to persistence *in vivo*. Comparative analysis with published proteins with antigenic properties makes it possible to distinguish immunoreactive proteins (40) among the proteins bearing a positive FC in dormancy, which may include specific antigens of latent tuberculosis. Additionally, the biotransformatory enzymes (oxidoreductases and hydrolases) capable of prodrug activation and stored up in the dormant state were annotated. These findings may potentially lead to the discovery of immunodiagnostic tests for early latent tuberculosis and trigger the discovery of efficient drugs/prodrugs with potency against non-replicating, dormant populations of mycobacteria.

## 1 Introduction

Tuberculosis (TB), caused by the pathogenic microorganism *Mycobacterium tuberculosis* (*Mtb*), currently surpasses HIV as the most common cause of mortality from an infectious disease worldwide (WHO global report).

In the course of infection establishment, *Mtb* passes through a series of intra- and extracellular locations – from alveolar macrophages to extracellular lesions, being exposed to various stress conditions including lowered pH values, nutrient and cofactor limitation, and peroxynitrite stress (1). Nonetheless, due to its unique metabolic plasticity, the bacterium survives these attacks, gradually transiting into a state of low metabolic activity – dormancy, associated with treatment failure for latent TB infection (2, 3). Bacterial internalization within the host is commanded by virulence factors on the one hand and reprogrammed metabolic networks, balancing the anabolic and catabolic processes with sufficient ATP generation to support survival (4–7).

Disseminated in the host, dormant bacilli may persist life-long; however, approx. 10% of latently infected TB carriers develop active disease (3). Dormant bacteria obtained from *in vitro* models are characterized by altered morphological traits (such as a thickened cell wall or cell-size diminishment), or resistance to antibiotics (8, 9). However, the cells obtained in such ‘quick’ models do not reflect mycobacterial physiology *in vivo*, where bacilli transit to a state of ‘non-culturability’ – a specific reverse state of inability to grow on solid media, and dependent on a reactivation procedure in selective liquid medium (10).

Whether metabolic pathways are active in such a state of ‘non-culturable’ dormancy of *Mtb* is not known and any knowledge in this respect will be valuable for understanding the survival mechanism and for selection and finding a remedy against latent TB. To elucidate such mechanisms, knowledge of the enzymes stored in long-term stored dormant cells is essential.

In the case of dormant cells that are metabolically inert, and therefore tolerant to common antibiotics, there is an attractive idea to consider ubiquitous enzymatic processes stored in dormancy as targets for the conversion of prodrug compounds into substances with unspecific antibacterial activity (like reactive oxygen and nitrogen species or antimetabolites) (11, 12).

Therefore, analysis of the enzymes stored up in dormancy may be prospective in terms of further development of follow-up prodrugs.

In the current study we analysed the proteomic profiles of *Mtb* cells in the dormant state (after 4 months of storage) in an *in vitro* dormancy model developed earlier (13). The advantages of this model over the widely exploited model of oxygen depletion (Wayne’s model (8)) comprise: the cells reveal a low metabolic activity (judging by the negligible rate of radiolabelled uracil incorporation, low level of transmembrane potential and undetectable respiration rate (14) as well as antibiotic tolerance); the cells demonstrate inability to grow on typical solid media without a procedure of reactivation – which is considered to be an intrinsic stage of genuine dormancy (15), developing therefore the state of ‘non-culturability’ (13).

In our recent publication, using 2D electrophoresis-based proteomics, we found that 1-year-old dormant, ‘non-culturable’ cells contain a significant amount of diverse proteins, including enzymes belonging to different biochemical pathways (16). However, this approach has evident limitations connected, firstly, with only partial coverage of the whole *Mtb* proteome and, secondly, with the inability to quantitatively estimate the differences in the abundancy of particular proteins between the groups of cells analysed (active vs dormant cells). LC-MS-based proteomic technologies are ideally suited for such comparative analysis, evading the above-mentioned problems of a 2D approach.

## 2 Materials and Methods

### 2.1 Bacterial strains, growth media and culture conditions

Inoculum was initially grown from frozen stock stored at −70 °C. *Mtb* strain H37Rv was grown for 8 days (up to OD600 = 2.0) in Middlebrook 7H9 liquid medium (HiMedia, India) supplemented with 0.05% Tween 80 and 10% ADC (albumin, dextrose, catalase) growth supplement (HiMedia, India). One millilitre of the initial culture was added to 200 mL of modified Sauton medium containing (per litre): KH_2_PO_4_, 0.5 g; MgSO_4_.7H_2_O, 1.4 g; L-asparagine, 4 g; glycerol, 2 mL; ferric ammonium citrate, 0.05 g; citric acid, 2 g; 1% ZnSO_4_.7H_2_O, 0.1 mL; pH 6.0–6.2 (adjusted with 1 M NaOH), and supplemented with 0.5% BSA (Cohn Analog, Sigma), 0.025% tyloxapol and 5% glucose. Cultures were incubated in 500 mL flasks containing 200 mL of modified Sauton medium at 37 °C with shaking at 200 rpm (Innova, New Brunswick) for 30–50 days, and pH values were periodically measured. In log phase, the pH of the culture reached 7.5–8.0 and then decreased in the stationary phase. When the medium in post-stationary phase *Mtb* cultures reached pH 6.0–6.2 (after 30–45 days of incubation), cultures (50 mL) were transferred to 50 mL plastic tightly capped tubes and kept under static conditions, without agitation, at room temperature for up to 5 months post-inoculation. At the time of transfer, 2-(N-morpholino)-ethane sulphonic acid (MES) was added at a final concentration of 100 mM to dormant cell cultures to prevent fast acidification of the spent medium during long-term storage.

#### 2.1.1 Evaluation of cell viability by CFU and MPN assays

Bacterial samples were serially diluted in fresh Sauton medium supplemented with 0.05% tyloxapol; three replicates of each sample from each dilution were spotted on agar plates (1.5%) with Middlebrook 7H9 medium (HiMedia, India), supplemented with 10% (v/v) ADC (HiMedia, India). Plates were incubated for 30 days at 37 °C, after which the number of CFU was counted.

The most probable number (MPN) assay is a statistical approach based on diluting a bacterial suspension until reaching the point when no cell is transited into any well. The growth pattern in three tubes is compared to the published statistical tables in order to estimate the total number of bacteria (either viable or dormant) present in the sample (17).

The MPN assay was carried out in 48-well plastic plates filled with 0.9 mL of special medium for the most effective reactivation of dormant *Mtb* cells. This medium contains 3.25 g of nutrient broth (HiMedia, India) dissolved in 1 L of a mixture of Sauton medium (0.5 g KH_2_PO_4_; 1.4 g MgSO_4_.7H_2_O; 4 g L-asparagine; 0.05 g ferric ammonium citrate; 2 g sodium citrate; 0.01% (w/v) ZnSO_4_.7H_2_O per litre, pH 7.0), Middlebrook 7H9 liquid medium (HiMedia, India) and RPMI (Thermo Fisher Scientific, USA) (1 : 1 : 1) supplemented with 0.5% v/v glycerol, 0.05% v/v Tween 80 and 10% ADC (HiMedia, India) (16). Appropriate serial dilutions of the cells (100 µL) were added to each well. Plates were sealed and incubated at 37 °C statically for 30 days. The MPN values, based on the well pattern originating from the visible bacterial growth at the corresponding dilution point, were calculated (17).

### 2.2 Preparation of samples for proteome analysis

Active and dormant cells were prepared in three biological replicates. Bacteria were harvested by centrifugation at 8000 *g* for 15 min and washed 10 times with a buffer containing (per litre) 8 g NaCl, 0.2 g KCl and 0.24 g Na_2_HPO_4_ (pH 7.4). The bacterial pellet was re-suspended in ice-cold 100 mM HEPES (4-(2-hydroxyethyl)-1-piperazineethanesulphonic acid) buffer (pH 8.0) containing complete protease inhibitor cocktail (Sigma, USA) and PMSF (phenylmethanesulphonyl fluoride) then disrupted with zirconium beads on a bead beater homogenizer (MP Biomedicals FastPrep-24) for 1 min, 5 times for active cells and 10 times for dormant cells. The bacterial lysate was centrifuged at 25,000 *g* for 15 min at 4 °C. To maximize protein isolation, SDS (2% w/v) extraction was carried out. The extracts were precipitated using a ReadyPrep 2D Cleanup kit (Bio-Rad, USA) to remove ionic contaminants such as detergents, lipids and phenolic compounds from protein samples.

### 2.3 In-solution digestion

The protein extracts were dissolved in 20 μL of 6 M urea in 0.1 M ammonium bicarbonate. The protein concentration was measured using a BCA Protein Assay Kit (Thermo Fisher Scientific, Waltham, MA, USA). An equal amount of the total proteins was used for further trypsinolysis and profiling.

Proteins were reduced using TCEP at 25 °C for 45 min, following alkylation with iodoacetamide (IAA) in the dark for 30 min, having final concentrations of 5 mM and 55 mM, respectively. Excess IAA was quenched with 1,4-dithiothreitol, then 50 mM ammonium bicarbonate was added to a total volume of 200 μL. Trypsin was added in a protein-to-trypsin ratio of 50 : 1, and tryptic digestion was done at 37 °C overnight. The reaction was stopped by adding formic acid (FA) to a final concentration of 5%. The peptide mixtures obtained were pre-fractionated and purified using a C18 Empore™ disk (3M, St. Paul, USA) as described in (18).

### 2.4 LC-MS/MS analyses

Prior to analysis, peptides were dissolved in 20 µL of 3% ACN/0.1% FA. The injection volume was 1 µL with a flow rate of 5 µL/min. For the mobile phases, 0.1% FA in MS-safe water (A) and 0.1% FA in 100% ACN (B) were used.

Analysis of peptides was done on a Synapt G2-Si High Definition Mass Spectrometer employing T-wave ion mobility powered mass spectrometry coupled to an ACQUITY UPLC M-Class System (Waters) using data-independent acquisition.

The UPLC system was equipped with a trapping column (nanoEase MZ Symmetry C18 Trap Column, 180 µm × 20 mm, 5.0 µm particle diameter, 100 Å pore size, Waters) and an analytical column (nanoEase MZ HSS T3 Column, 75 µm × 100 mm, C18, 1.8 µm particle diameter, 100 Å pore size, Waters). The sample was loaded onto the trapping column for 2 min and transferred to the analytical column. The peptides were eluted from the analytical column with a flow rate of 0.4 μL/min with a linear gradient of increasing concentration of mobile phase B from 5% to 35% for 70 min. Within the nanoFlow^TM^ ESI-source, a capillary voltage of 2.5 kV was used, the sampling cone voltage was 40 V and the source offset was 80 V. The source temperature was set to 80 °C. In the trapping, ion mobility and transfer chambers, the wave velocity of the TriWave was set to 1000, 650 and 175 m/s, respectively, and ramping wave heights of 40, 40 and 4.0 V were used, respectively. In the HDMS^E^ mode, the collision energy was ramped to 22–45 eV. Data acquisition was carried out using MassLynx software (Waters). HDMS^E^ mass data were acquired in low and high energy modes. Raw data were noise-reduced using the Noise Compression Tool (Waters).

The acquired data were submitted for processing and database searching using Progenesis software against the database prepared in-house containing protein sequences from *Mycobacterium tuberculosis* strain ATCC 25618/H37Rv downloaded from the UniProt database (version 20190304) supplemented with sequences of common contaminants (Max Planck Institute of Biochemistry, Martinsried, Germany). The low-energy and high-energy threshold values were set to 150 and 50 counts, respectively. The parameters used for the database search were: enzyme specificity: trypsin, allowed missed cleavages: 2, fixed modification: carbamidomethylation (Cys), variable modifications: N-terminal protein acetylation, and oxidation (Met); the false discovery rate was 1%; minimum fragment ion matches per peptide, minimum fragment ion matches per protein and minimum peptides per protein were set to 1, 4 and 2, respectively. Proteins were considered as significant if they were identified in at least two independent biological replicates.

### 2.5 Statistical analysis

Data analysis and visualization were carried out in the R environment, applying scripts developed in-house.

#### 2.5.1 Data normalization

Before the analysis, the total protein concentration was quantified and an equal amount of protein extracts was used for further trypsinolysis and profiling – pre-instrumental normalization on total protein concentration was carried out (19). To make the profiles comparable for the subsequent statistical analysis, post-instrumental pre-treatment methods were carried out: to deal with heteroscedasticity and to make skewed distributions symmetric, *log_2_* transformation was performed. To force the samples to have the same distributions for the protein intensities, quantile normalization was carried out (20).

#### 2.5.2 Principle component analysis and hierarchical clustering

For illustration of intragroup and between-group variances, principle component analysis (PCA) was carried out, applying the preliminarily normalized, *log_2_*-transformed and Pareto-scaled datasets (21–23). Similarity/dissimilarity between the groups was determined by a method of hierarchical clustering using the method of Ward (24).

#### 2.5.3 Z-score calculations and heatmap visualization

A pre-treated protein matrix, consisting of proteins arranged in rows and groups of samples arranged into columns, was centred and scaled (25). The centring represents the difference between the value of a protein’s abundancy and the total mean of the protein. The scaling is the result of division of the scaled metabolite by its σ. The resulting Z-scores, being in a similar range for the palette of proteins, make feasible comparison of the variances between the variables (proteins).

#### 2.5.4 Statistical significance analysis

The corresponding *p*-values of comparison of the two groups of cells (dormant vs active) were calculated from a two-sided *t*-test. Since the data matrix was initially *log_2_*-transformed, the resulting *log_2_(FC_dorm/mean_)* is the difference between the *log_2_(mean_dorm_)* and *log_2_(mean_act_)*.

### 2.6 Subtractive proteomic analysis of mycobacterial proteins prevailing in dormancy with antigenic properties

The basic principles of subtractive analysis developed previously were used in the current work [26]. The significantly changed proteins revealing in dormancy a positive fold change (FC) were subjected to subtractive analysis. In the first step, the paralogue proteins were identified using a CD-HIT online platform (28, 29). The proteins with 60% identity were considered to be paralogous and discarded from the further analytical steps. In the next step, the BLASTp procedure was applied to select non-homologues to *Homo sapiens* proteins in the query set, using a threshold expectation value of 0.001 and similarity below 35%. The resulting list of proteins was further categorized according to their subcellular localization using the CELLO online platform (30).

The shortened list of proteins was further compared with previously published datasets with annotated *Mtb* proteins with antigenic properties: i) a list of 181 proteins detected exclusively in the sera of TB-positive patients from a study on the dynamic antibody response to the proteome (31); ii) a list of 62 proteins resulting from serodiagnosis of latent *Mtb* infection (32); iii) a list of 22 proteins resulting from the analysis of TB-specific IgG antibody profiles, detectable on two different platforms (Luminex and MBio) (33).

### 2.7 Annotation of the list of enzymes – prodrug activators

KEGG API was used to prescribe EC numbers to the initially prescribed UniProt KO code (34).

## 3 Results

### 3.1 Dormant mycobacterial cells subjected to the proteomic analysis and the results of proteomic profiling

Dormant *Mtb* cells subjected to the further proteomic profiling bore the typical traits of ‘non-culturability’ (13, 16), particularly incapability to grow on rich solid medium: the experimentally estimated viability by the CFU method was 1.32 ± 0.48 × 10^4^ cells/mL (± SE) for dormant cells and 1.17 ± 0.44 × 10^8^ cells/mL (± SE) for active cells. At the same time, estimation of the total number of viable but ‘non-culturable’ cells after resuscitation in fresh liquid medium by the MPN assay resulted in 1.21 ± 0.65 × 10^9^ cells/mL (± SE) for dormant cells and 0.97 ± 0.44 × 10^9^ cells/mL (± SE) for actively growing cells (Fig. 1). Before the analysis, the total protein concentration was quantified as 3.56 ± 0.24 mg/g wet weight (± SE) for dormant cells and 6.01 ± 0.14 mg/g wet weight (± SE) for active cells. An equal amount of the total proteins was used for further trypsinolysis and LC-MS/MS profiling (19). LC-MS profiling resulted in the detection of 1379 proteins (whose encoding genes Rv were further used as identifiers), covering 35% of the coding *M. tuberculosis* H37Rv genome. However, the raw instrumental data showed skewed distributions of protein intensity, tending to lower values (Fig. 2A).

**Fig. 1.**
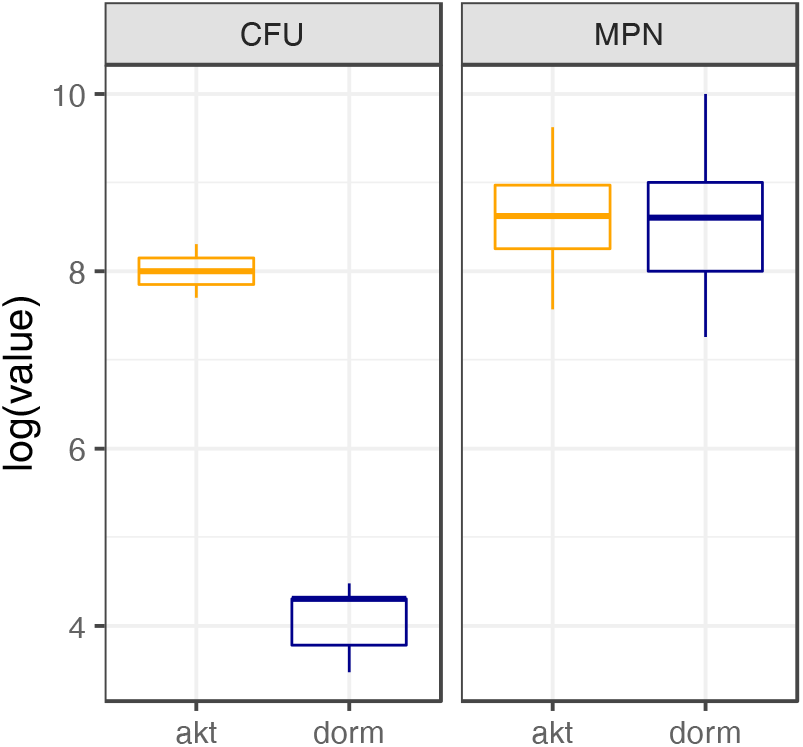
Viability assays (CFU and MPN) of *M. tuberculosis* H37Rv cells subjected to further proteomic analysis. Active cells were probed in the late log phase – the 10^th^ day of cultivation. Viability of the dormant cells was analysed after 4 months of storage after gradual acidification of the bacterial culture in the post-stationary phase (81). Estimation based on probing of 6–9 independent samples of each type of cell.

**Fig. 2.**
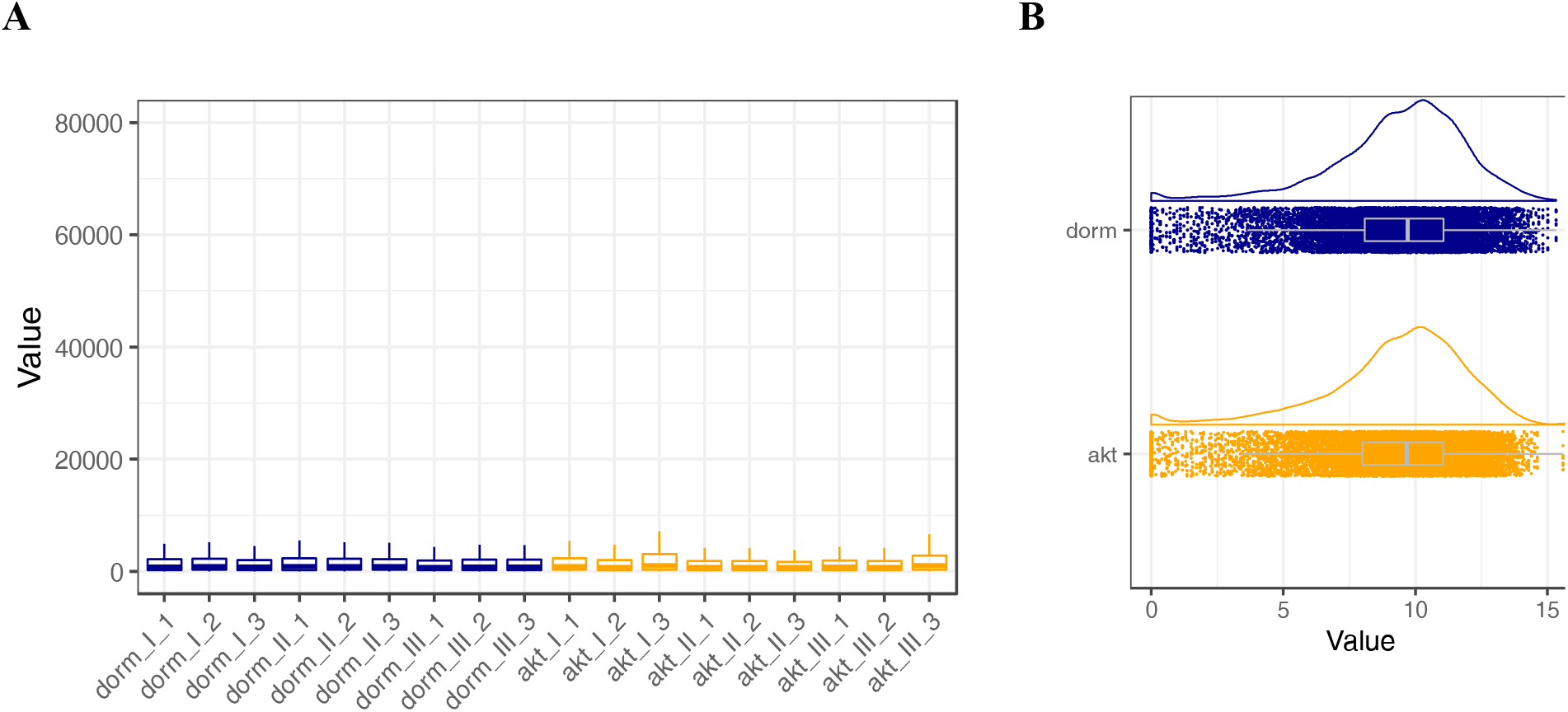
Normalization of proteomic profiles. A) Distribution of raw protein intensities; B) average median intensity of nine samples (biological and technical replicates) for each physiological condition after *log_2_* transformation and quantile normalization of the protein profiles.

For the subsequent statistical analysis, post-instrumental pre-treatment methods were carried out: to deal with heteroscedasticity and to make skewed distributions symmetric, *log_2_*-transformation was performed. To force the samples to have the same distributions for the protein intensities, quantile normalization was carried out (20), resulting in comparable normally distributed profiles with equal medians, making the protein profiles eligible for further statistical analyses (Fig. 2B).

### 3.2 PCA and hierarchical clustering

The results of PCA (23) depict ca. 80% of all variance in the protein profiles. To perform PCA, the data matrix was preliminary Pareto-scaled (22). The results demonstrate that samples are clearly distinguished by their physiological state (‘dormant’ vs ‘active’) (Fig. 3). Hierarchical cluster analysis, based on calculations of the amount of within-cluster dissimilarity analogously results in two distinct clusters, separating dormant cells from active ones. Ward’s minimum variance method was used for the calculations (24) (Fig. 3B).

**Fig. 3.**
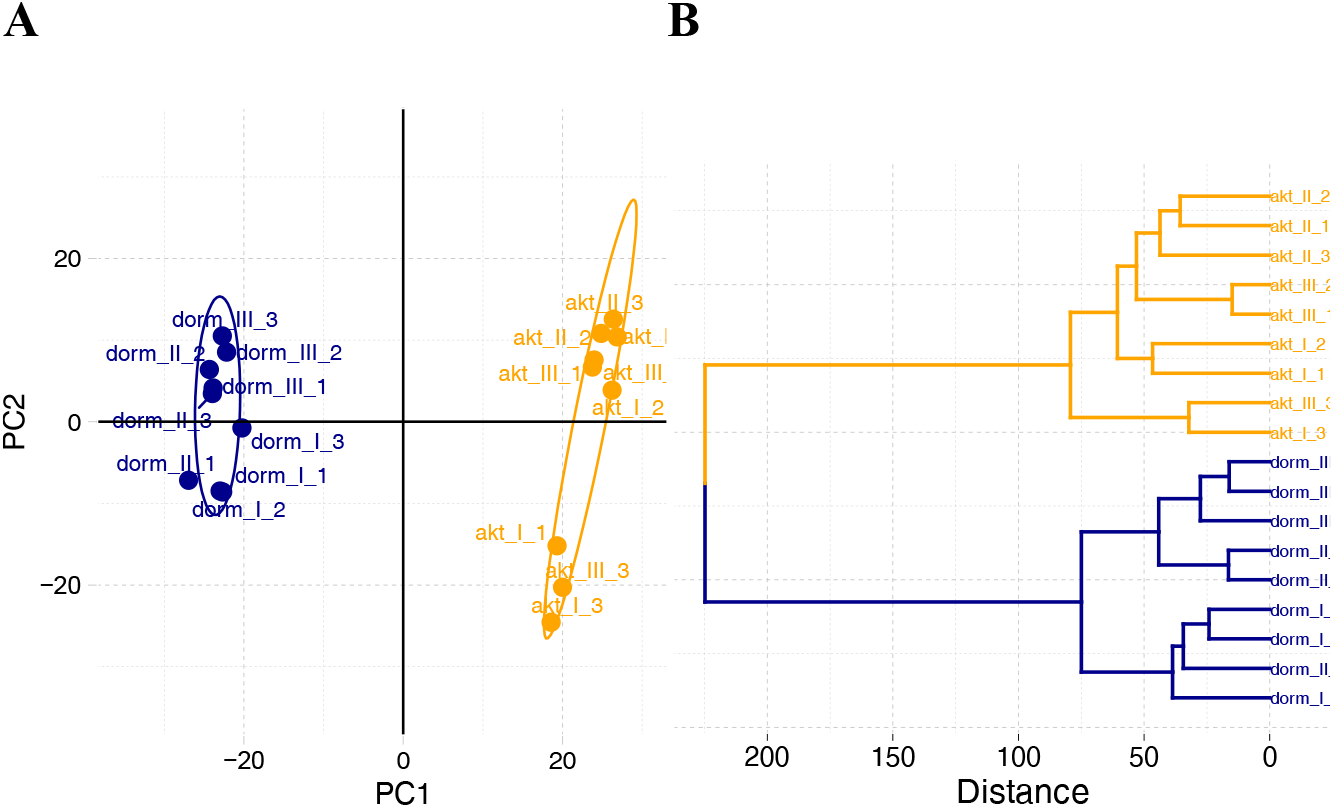
Results of the principal component analysis (A) and hierarchical clustering (B) of *M. tuberculosis* cells used for proteomic profiling. PC1 with the largest variance of each protein level separates the samples into two distinct clusters. The elliptical boundary is analogous to the 95% CI for both bivariate distributions. Ward’s method (24) of hierarchical clustering similarly demonstrates segregation of the two types of physiological group.

### 3.3 Differential analysis

The results of differential analysis are listed in the Supplement and graphically depicted as a volcano plot in Fig. 4A. Generally, the abundancy of 894 proteins (out of 1379) was found to change significantly (*p* < 0.05) in the transition to the dormant state – among them 468 with a positive FC (‘stored in dormancy proteins’) and 426 with a negative FC (‘decreased in dormancy proteins’). The rest of the proteins (485) lay beyond the scope of differential analysis – there was greater variance in their fluctuation in transition to dormancy and consequently their changes were insignificant. Analysis of the first 30 proteins stored significantly differently in dormancy, bearing the maximum *log_2_FC*, discloses a cohort of regulatory proteins and virulence factors (and not enzymatic proteins, dictating biochemical adaptation to the stress conditions), despite the *in vitro* nature of the dormancy model (Fig. 4B). According to the main stress factor – pH decrease – we observed upregulation of the transcriptional regulator phoP (Rv0757, *p* < 0.05, *log_2_FC* = 1.43) of the two-component phoP–phoR system, conferring the fitness of mycobacteria to the lowered pH values [16,35,36]. In addition, phoPR controls secretion of the antigens Ag85A (Rv3804c, *p* < 0.05, *log_2_FC* = 1.28) and Ag85B (Rv0129c, *p* > 0.05, *log_2_FC* = 1.11) as well as ESAT-6 EsxA (Rv3875, *p* < 0.05, *log_2_FC* = 4.14) – a secreted strong human T-cell antigen protein that plays a number of roles in modulating the host’s immune response to infection (37) (Fig. 4C). Tightly associated with the ESX-1 and ESX-5 secretion systems are proteins of the PE/PPE family (38). Out of 95 KEGG-annotated PE/PPE proteins we detected six: PPE18 (Rv1196, *p* < 0.05, *log_2_FC* = 2.44), PPE19 (Rv1361c, *p* < 0.05, *log_2_FC* = 2.11), PPE32 (Rv1808, *p* < 0.05, *log_2_FC* = 3.15), PPE60 (Rv3478, *p* < 0.05, *log_2_FC* = 1.05), PE15 (Rv1386, *p* < 0.05, *log_2_FC* = 3.64) and PE31 (Rv3477, *p* > 0.05, *log_2_FC* = 0.39). The low discovery rate may be explained by the paucity of trypsin-cleavage sites in these proteins and their structural homology (39). For the proteins of this class, a variety of functions is assumed: many PE and PPE genes have been shown to be upregulated in starvation and during stress conditions (40). A chaperone function of PE proteins has been reported, particularly in transporting of MPT64 antigen protein (Rv1980c, *p* < 0.05, *log_2_FC* = 0.69) (41). Additionally, the phoP transcriptional regulator has been shown to downregulate devR (Rv3133c, *p* > 0.05, *log_2_FC* = 0.01), a regulator of the DosR ‘dormancy’ regulon (35, 42). Correspondingly, out of ca. 50 postulated genes of the DosR regulon, only 19 proteins were detected in the current study, 14 of which were found significantly stored in the dormant state (Fig. 4D). Similarly, we observed a down-shift of a number of other transcriptional regulatory proteins, e.g., Rv0275c, Rv1556, Rv1019, Rv0818, Rv3058c, Rv3249c, etc. Analogously, in the absence of translational processes, the abundancy of translation initiation factors (Rv3462c, Rv2839c) as well as 30s and 50s ribosomal proteins (Rv0717, Rv2875c, Rv0641) was similarly decreased.

**Fig. 4.**
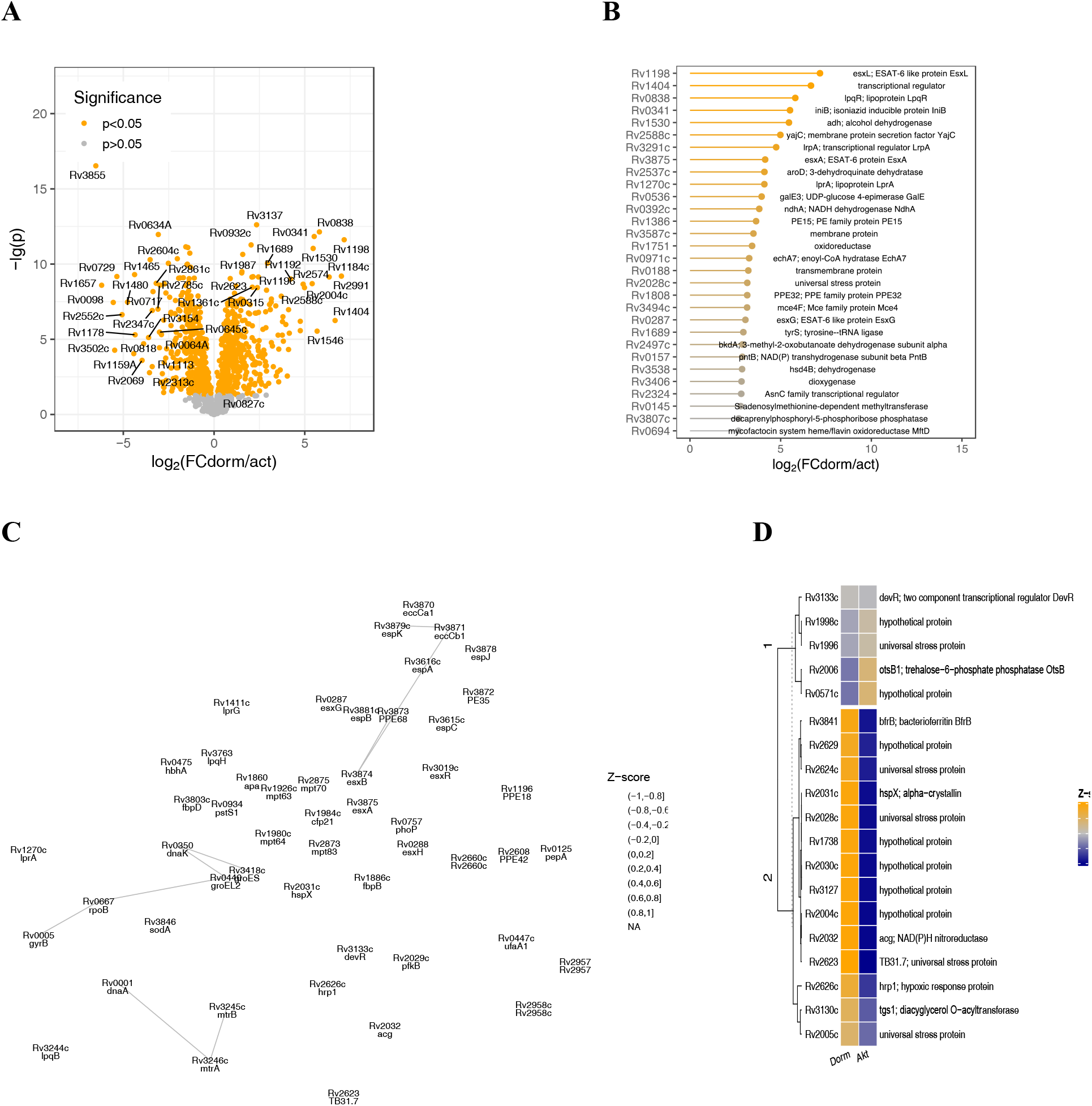
General statistical results of comparative proteomic analysis of dormant vs active *M. tuberculosis* cells. A) Results of differential comparison of dormant cells vs active cells reveal 894 statistically significantly changed proteins, among them 468 proteins prevailing in dormancy (with a positive FC value) and 426 proteins whose abundancy decreased in the transition to dormancy (with a negative FC value); B) visualization of the 30 proteins most abundant in dormancy with the maximum FC; C) STRING-based interaction network of phoP and devR regulators of DosR regulon. Upregulation of pH-dependent regulator phoP suppresses devR, a regulator of the DosR system, however activating the expression of mycobacterial virulence factors: ESAT and PE/PPE proteins; D) heat map of the detected proteins of DosR regulon (14 upregulated out of 50 postulated for Wayne’s model of dormancy) (82).

There was no consistency in behaviour of proteins of the ABC trehalose transporter complex LpqY– SugA–SugB–SugC (Rv1235, Rv1236, Rv1237, Rv1238): out of four annotated in the *Mtb* genome only Rv1235 and Rv1238 were detected in the current study, albeit with a different sign of FC in dormancy, rather indicating a decrease in their abundancy and efficacy in the current stress conditions and confirming the dependency of the dormant cells on functioning trehalose to replenish intracellular pathways (43). On the contrary, the components of the phosphate ABC transporter pstS1-3 (Rv0928, Rv0934, Rv0932c) were found concordantly to be stored in dormancy, indicating a possible dependency of dormant cells on extracellular sources of inorganic phosphate. Part of the ABC efflux pump Rv1218c, conferring the cells the resistance to a wide range of drugs, was found to be similarly stored.

For the analysis of biochemical adaptation to the transition to dormancy, the 666 detected enzymes (both with positive and negative *log_2_FC*, based on KEGG orthology annotation) were placed on the metabolic map and the corresponding processes which they catalyse were highlighted: in orange for those enzymes with a positive FC (enzymes stored in dormancy) and in blue for those demonstrating a negative FC (a relative decrease in dormancy). The width of the highlighted lines corresponds to the *p*-value (the thicker lines are for *p* < 0.01, the thinnest lines are for insignificant results (*p* > 0.05)) (Fig. 5A). A glance at the general metabolic visualization (Fig. 5A) gives an impression that, in dormancy, a scattered ‘mosaic’ intactness of enzymes can be seen: the flawless functionality of the many pathways is hindered by enzymes significantly decreased in dormancy, e.g., the functionality of the urea cycle may be broken because of the decreased level of argininosuccinate synthase (Rv1658, *p* < 0.05, *log_2_FC* = −0.69). To annotate precisely the metabolic pathways differentially preserved in dormancy, pathway enrichment analysis was carried out on the enzymatic proteins with FC > 0 and *p* < 0.05 (44, 45) (Fig. 5B). Detailed pathway analysis revealed nine upregulated proteins of TCA; however, isocitrate dehydrogenase (Rv0066c, *p* < 0.05, *log_2_FC* = −0.41) and subunits of α-ketoglutarate dehydrogenase (Rv2454 and Rv2455) demonstrated negative values of *log_2_FC*, impeding the normal flow of the TCA cycle (Fig. 5C). The subunits of membrane-bound fumarate reductase were not technically detected; however, subunits of the succinate dehydrogenase complex (Rv3318 and Rv0247c) were detected, potentially enabling the reverse flow of the processes in the ‘reductive branch’ of the TCA cycle (part of the Arnon–Buchanan cycle) from pyruvate to succinate, providing the cell with reoxidized NAD^+^. The intactness of enzymes acting in the β-oxidation of fatty acids may provide the cell with a surplus of acetyl-CoA (in the case of oxidation of even-chained fatty acids) and propionyl-CoA (in the case of oxidation of odd-chained fatty acids). Dormant *Mtb* cells demonstrated the presence of enzymes of the methylcitrate cycle, which may allow the conversion of propionyl-CoA and oxaloacetate to succinate and pyruvate. The latter can be converted to fumarate by the detected succinate dehydrogenase (Rv3318, Rv0247c). If this pathway is functioning, the accumulation of organic acids (contributing to sustainment of the energized level of transmembrane potential in dormancy (46, 47)) in the cells and medium should be expected; that, however, was found experimentally for dormant cells of *M. smegmatis*, obtained in the same model of self-acidification of growth medium (14). Contributing to the accumulation of these acids and to detoxification of surplus acetyl-CoA is the glyoxylate pathway, whose enzymes isocitrate lyase (Rv0467) and malate synthase (Rv1837c) were found to be accumulated in dormant cells.

**Fig. 5.**
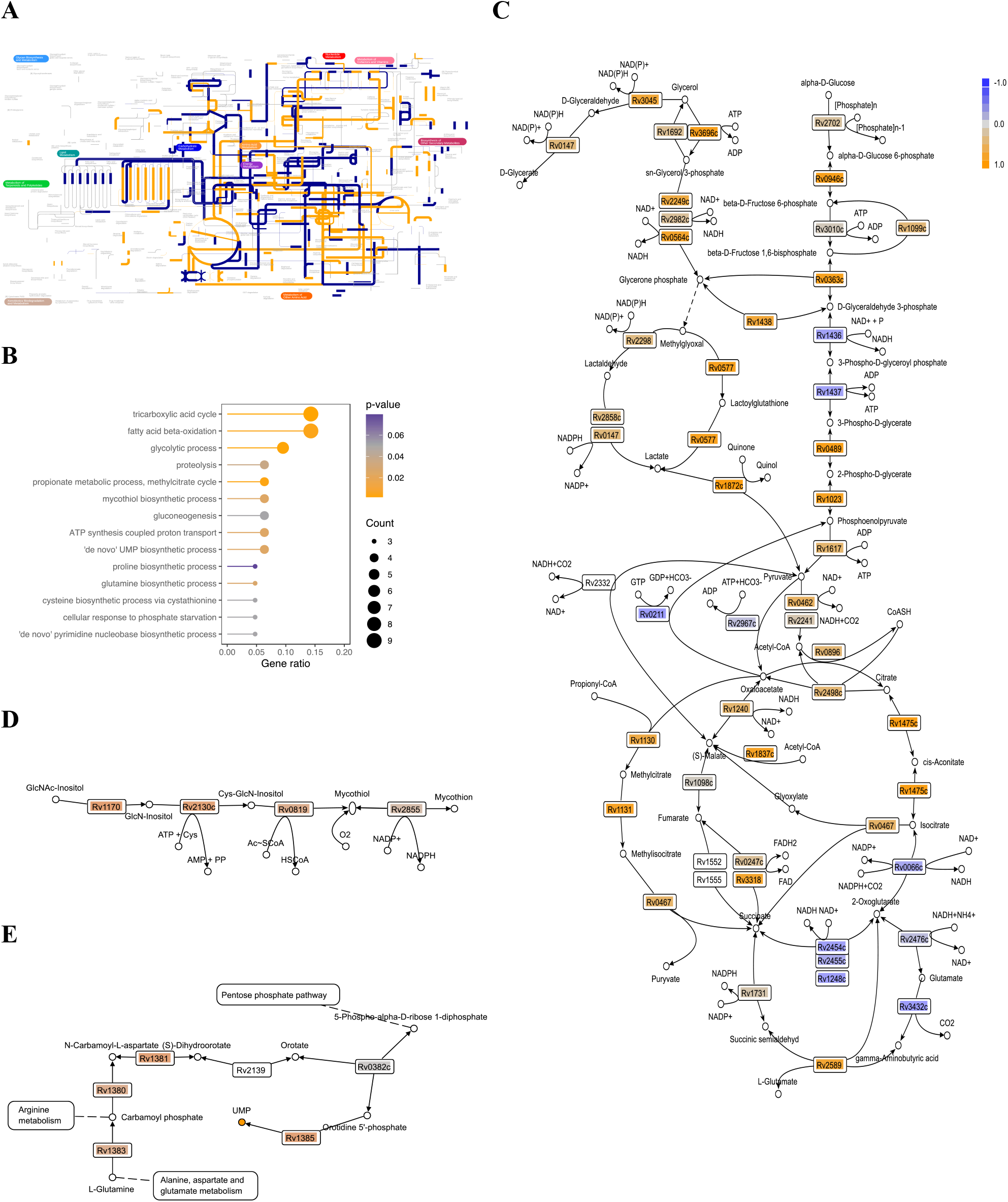
Proteomic composition of enzymatic proteins in dormant *M. tuberculosis*. A) Pathway visualization, based on KO annotation: the detected enzymes (either stored in the dormant state, or those whose abundancy decreased) were placed on a metabolic map and the corresponding processes highlighted – orange for enzymes stored in dormancy (FC > 0), dark blue for degraded enzymes (FC < 0); line thickness corresponds to statistical significance – the thickest lines depict statistically valuable processes; B) GO enrichment analysis for the pathways assembled by the enzymes stored in dormancy; C) reconstruction of central carbon metabolism pathways of dormant *M. tuberculosis* cells – Z-scores are mapped on the corresponding Rv coding enzymes; D) mycothiol salvage pathway; E) reconstruction of ‘*de novo*’ UMP pathway. UMP accumulation in the dormant state was similarly recently observed in the same model conditions in *M. smegmatis* cells (14).

The classical flow of the glycolytic pathway is hampered by deficient glyceraldehyde-3-phosphate dehydrogenase (Rv1436, *p* < 0.05, *log_2_FC* = −0.43) and phosphoglycerate kinase (Rv1437, *p* < 0.05, *log_2_FC* = −0.61); however, there is a bypass, consisting of the formation of glycerone phosphate, which can further spontaneously convert into methylglyoxal, which in further detoxification pathways leads to pyruvate through lactate formation. Similarly, all enzymes of glycerol metabolism were detected – the catabolism of glycerol or glycerol-3-phosphate is conjugated with the toxic carbonyl compounds methylglyoxal and lactaldehyde (the latter was detected experimentally for *M. smegmatis* (14)). However, functioning aldehyde dehydrogenases (Rv0147, Rv2858c) or glyoxylase (Rv0577) detoxify these intermediates, ultimately to lactate. The further transformation of forming lactate by a quinone-dependent lactate dehydrogenase finally supplies the cells with pyruvate capable of further catabolism in the TCA, glyoxylate and methylcitrate pathways. *Mtb* used to have annotated two lactate dehydrogenases, *lldD2* (Rv1872c, *p* < 0.05, *log_2_FC* = 1.04) and *lldD1* (Rv0694, *p* < 0.05, *log_2_FC* = 2.67); however, lactate dehydrogenase activity was confirmed for *lldD2* only (48). On the other hand, upregulation of Rv0694 was observed in the current conditions of dormancy in glycerol-glucose medium. Rv0694 is a member of the Rv0691–Rv0694 gene cluster, involved in biosynthesis of a recently discovered peptide-derived electron carrier – mycofactocin (49).

The decreased detection of particular enzymes may be due to the preservation of proteolytic peptidases abundant in the dormant proteome: Rv0319 (*p* < 0.05, *log_2_FC* = 1.1), Rv0457c (*p* < 0.05, *log_2_FC* = 0.82), Rv3883c (*p* < 0.05, *log_2_FC* = 2.53) and Rv0983 (*p* < 0.05, *log_2_FC* = 0.64).

The enzymes of glycerol catabolism that are stored unprocessed in dormancy contribute to lipid metabolism and may serve as a source of reoxidation of reductive equivalents (Fig. 5C). The gapped glycolytic pathway fails to serve substrate-level ATP generation (if any): however, accumulated polyphosphate (the transport of inorganic phosphate remains intact) may serve a phosphagen – a reservoir of phosphoryl groups, that can be used to generate ATP, thus we detected a reversible ppk1 (Rv2984, *p* < 0.05, *log_2_FC* = 0.57).

Additionally, we were able to reconstruct a *de novo* unimpaired uridine monophosphate (UMP) pathway, serving the precursor of all pyrimidine nucleotides.

Similarly to the known involvement of protective redox systems as a response to ROS attacks in macrophages, upregulation of superoxide dismutase was observed in our dormancy conditions (sodA, Rv3846, *p* < 0.05, *log_2_FC* = 1.04). Since mycobacteria lack glutathione, the ubiquitous low-molecular weight thiol in other organisms, *Mtb* is dependent on mycothiol for antioxidant activity; the enzymes encoding its biosynthesis were found to be stored in the dormant state (Fig. 5).

To the oxidative stress response system belong F_420_-dependent glucose-6-phosphate dehydrogenase *fgd1* (Rv0407, *p* > 0.05, *log_2_FC* = 0.71) and F_420_H_2_-dependent menaquinone reductase Rv1261c (*p* > 0.05, *log_2_FC* = 0.52), contributing to the protection of bacteria from the formation of toxic semiquinones (50). In shaping the response to ROS and RNS, the accumulation of the regulator sigH was observed (Rv3223c, *p* < 0.05, *log_2_FC* = 0.48).

### 3.4 Annotation of immunogenic proteins accumulated in dormant cells

It was interesting to search for immunogenic proteins among detected proteins with a positive FC in dormant *Mtb*, as they are potentially applicable for latent TB diagnosis. In the current study we enriched the possible candidates, based on the subtractive proteomic concept [26]. The essence of this approach presumes the identification of non-paralogue proteins followed by the identification of non-homologues to *H. sapiens* proteins and the analysis of subcellular localization (Fig. 6A). Thus, the initially applied 468 proteins significantly represented in dormancy with a positive FC resulted in 273 non-homologue proteins after the first two filters, further localization analysis of which allowed the identification of 53 membrane proteins, 175 cytoplasmic proteins and 45 extracellular proteins.

**Fig. 6.**
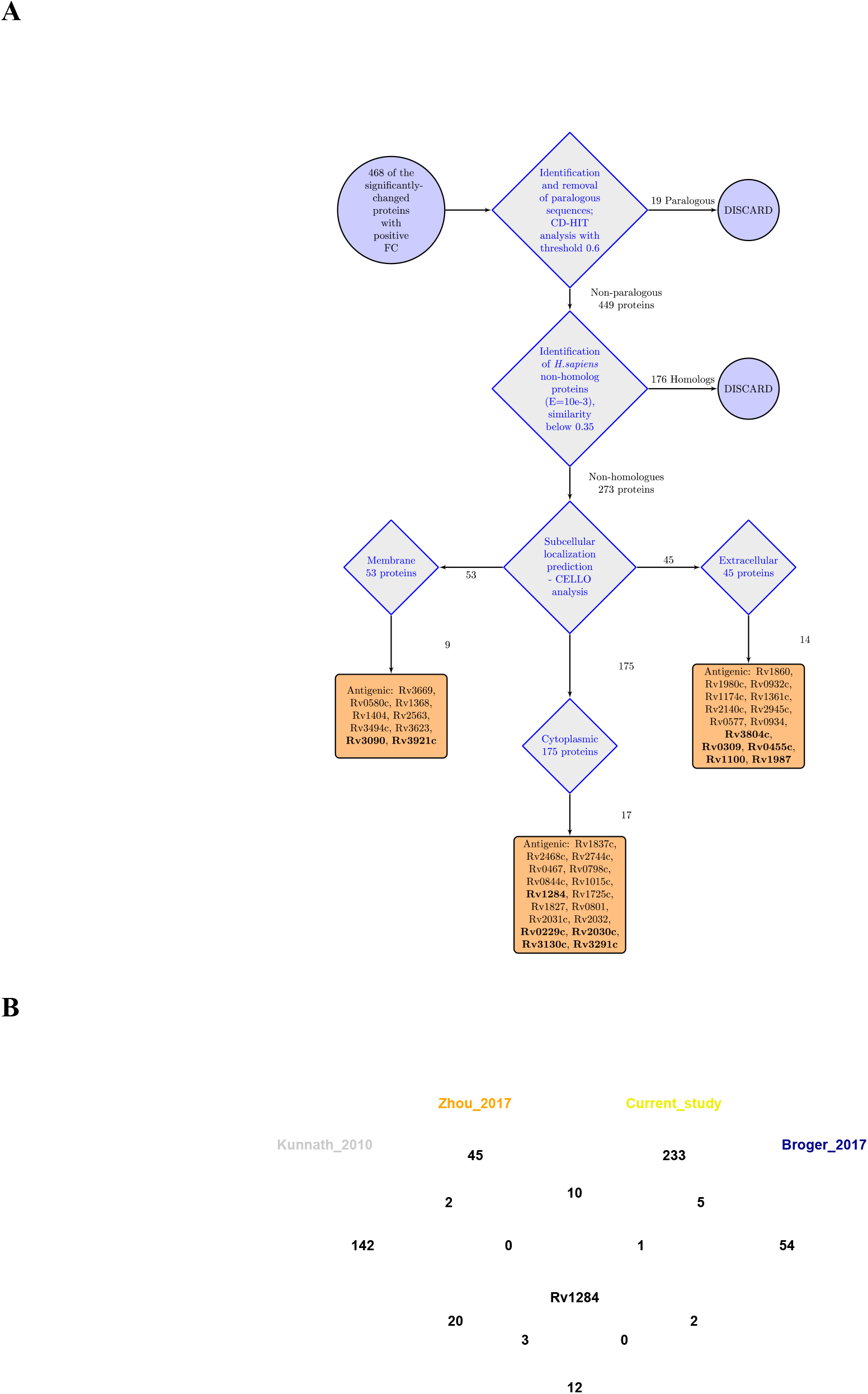
Determination of potential antigenic proteins in dormant *M. tuberculosis* proteome. A) General workflow for establishment of potential antigenic proteins. The complete list of antigenic proteins is available in the supplementary file; bold-annotated proteins are those with immunogenic properties selected by the sera of latently infected patients (32); B) Venn analysis of the reference antigenic proteins annotated from previously published works (31–33), with the list of proteins resulting from the subtractive proteomic analysis of the current data set.

The short-listed data set was compared with the reference lists from published works on studies of the sera of human TB patients (31–33).

The comparative analysis reference lists comprised the most reactive proteins (in total 265 antigens) as described in the Material and Methods.

Comparative analysis of the proteins in these reference sets with the selected proteins in the current study with a positive FC in dormancy revealed 40 potentially immunogenic proteins specific for dormancy (Fig. 6A). Remarkably, these studies (including our data set) demonstrate only one shared protein, Rv1284 – a carbonic anhydrase (*p* < 0.05, *log_2_FC* = 0.38), which was previously annotated among the latency-related antigens of *Mtb* (51) (Fig. 6B).

### 3.5 Annotation of proteins accumulated in dormancy – potential prodrug activators

The accumulation of a significant number of enzymatic proteins in the dormant state may imply the potency of ‘non-culturable’ *Mtb* cells to perform biotransformation reactions.

Analysis of the distribution of enzyme classes in dormancy revealed the preservation of a bulk bundle of oxidoreductases and hydrolases (Fig. 7A). The list of oxidoreductases (73) demonstrated in dormancy with an FC > 0 is summarized in the Supplement. The majority of the annotated enzymes of this class are NAD- and FAD-dependent, bearing relatively low standard reduction potentials (−0.32 and −0.22 V correspondingly). There are, however, two F_420_-dependent enzymes, Rv0407 (F_420_-dependent glucose-6-phosphate dehydrogenase Fgd1) and deazaflavin (F_420_)-dependent nitroreductase (Ddn), to detect. F_420_-dependent enzymes are known to bear a lower reduction potential (−0.32 V) and have been shown to participate in bicyclic nitroimidazole compounds (50, 52). Similarly, the FAD-dependent decaprenylphosphoryl-beta-D-ribose-2’-epimerase (catalysing the conversion of decaprenyl-phosphoryl-D-ribose to decaprenyl-phosphoryl-D-arabinose – the sole arabinose precursor in the pathway of arabinogalactan biosynthesis) is capable of nitroreducing recently developed benzothiazinones (BTZ) (53).

**Fig. 7.**
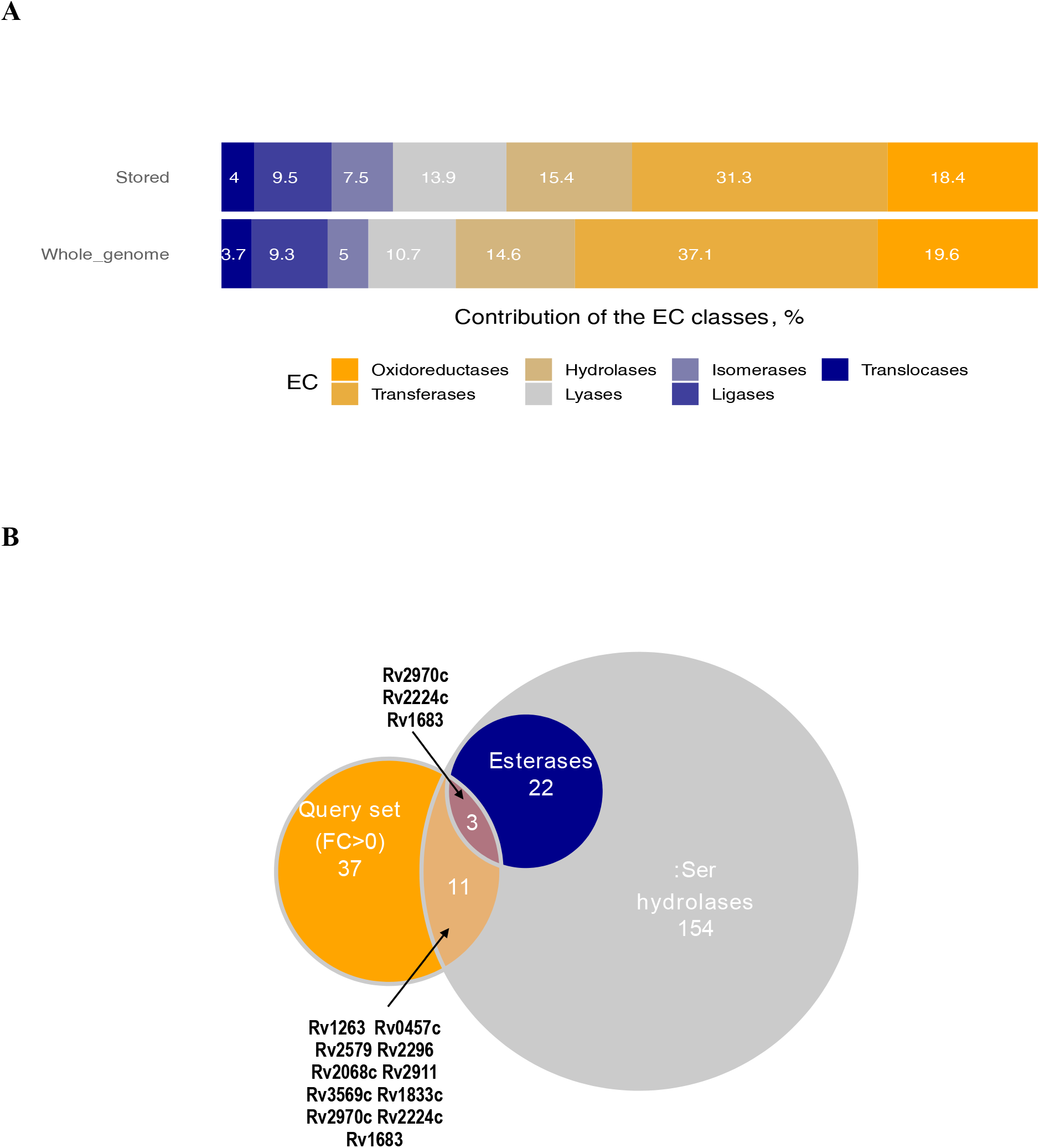
Enzymes of dormancy as potential intracellular prodrugs activators. A) Distribution of the enzyme classes within the enzymes stored in dormancy compared to whole-genome enzyme classes; B) overlap of the detected 48 hydrolytic enzymes with the annotated group of serine hydrolases and esterases (57).

The component of the non-proton pumping NADH:quinone oxidoreductase (Rv0392c, *p* < 0.05, *log_2_FC* = 3.81) accumulated in dormancy is hypothesized to reduce clofazimine, an antibiotic whose mode of action is mediated through ROS formation (54).

Hydrolytic enzymes (esterases and lipases) are considered to play a crucial role in mycobacterial persistence (55, 56). Among the enzymes detected with a positive FC in the dormant state, 48 could be classified as hydrolytic enzymes (Supplement). This list was compared with a list of annotated serine hydrolases and esterases (57) (Fig. 7B). In the query data set of enzymes with a positive FC, three enzymes potentially keep their activity in dormancy (Rv2970c, Rv2224c and Rv1683) (57). The list of enzymes of other EC classes can be found in the Supplement.

## 4 Discussion

The observation that the majority of proteins in dormancy are significantly abundant (in comparison to the active state), and bear a positive FC, corresponds to the previously reported observation on the ability of *Mtb* to preserve protein diversity in the dormant state (16). Comparison of the data originating from 2D electrophoresis analysis of D1 cells (4.5 months of dormancy) (16) with the same gene set from the current study (262 genes) after a common ranking procedure, followed by the Mann–Whitney test, reveals similarity between the data sets (*H_o_*: *U_LC-MS_* = *U_2D_*, *p*-value = 0.8894) (Fig. 7). Similar to the previously published 2D proteome analysis, only a limited number of accumulated proteins belong to the annotated DOS regulon, which demonstrates the differences between the anoxic Wayne’s model and the current model of self-acidification in the post-stationary phase. Analogously, the increased abundancy of Rv0757 – a component of the pH-dependent regulator phoP which is upregulated in D1 and D2 dormancy states – may contribute to the low discovery rate of proteins of the DOS system (16).

phoP is a virulence regulator which is conserved in a wide range of bacteria and controls the transcription of more than 600 genes (58). Besides its function of regulating the transcription of virulence factors (see the results of differential analysis - §3.3), it has been hypothesized that *phoP* coordinates gene expression with changes in membrane potential, and encodes the proteins that control the state of membrane energization (58). In *Salmonella*, *phoP* has been shown to control the production of superoxide dismutase, SOD (59), which was found to be abundant in dormancy in the current study. Correspondingly, the protective systems were detected in the current study – accumulation of SODs (sodA – Rv3846, sodC – Rv0432) and the DNA-binding proteins hupB (Rv2986) and iniB (Rv0341), as well as several chaperone proteins (DnaK – Rv0350, dnaJ1 – Rv0352). However, 2D analysis, being qualitative in its nature, could not disclose the changes in quantitative abundance of these proteins. Thus, e.g., catalase G (katG, Rv1908c) could be detected in D2 (16), the abundancy of which in the dormant state in the current study, however, was slightly decreased in comparison to the active state.

Moreover, the main drawback of the 2D approach is its inability to reach a meaningful coverage of biochemical processes – thus, GO enrichment analysis based on the enzymes detected in D1 (16) revealed only three statistically feasible pathways. Manual reconstruction of the glycolysis and TCA pathways demonstrated only a limited coverage of these pathways. Moreover, those remaining undisclosed were the transformation of glycerol, propionyl-CoA metabolism and the glyoxylate pathway (Fig. 8).

**Fig. 8.**
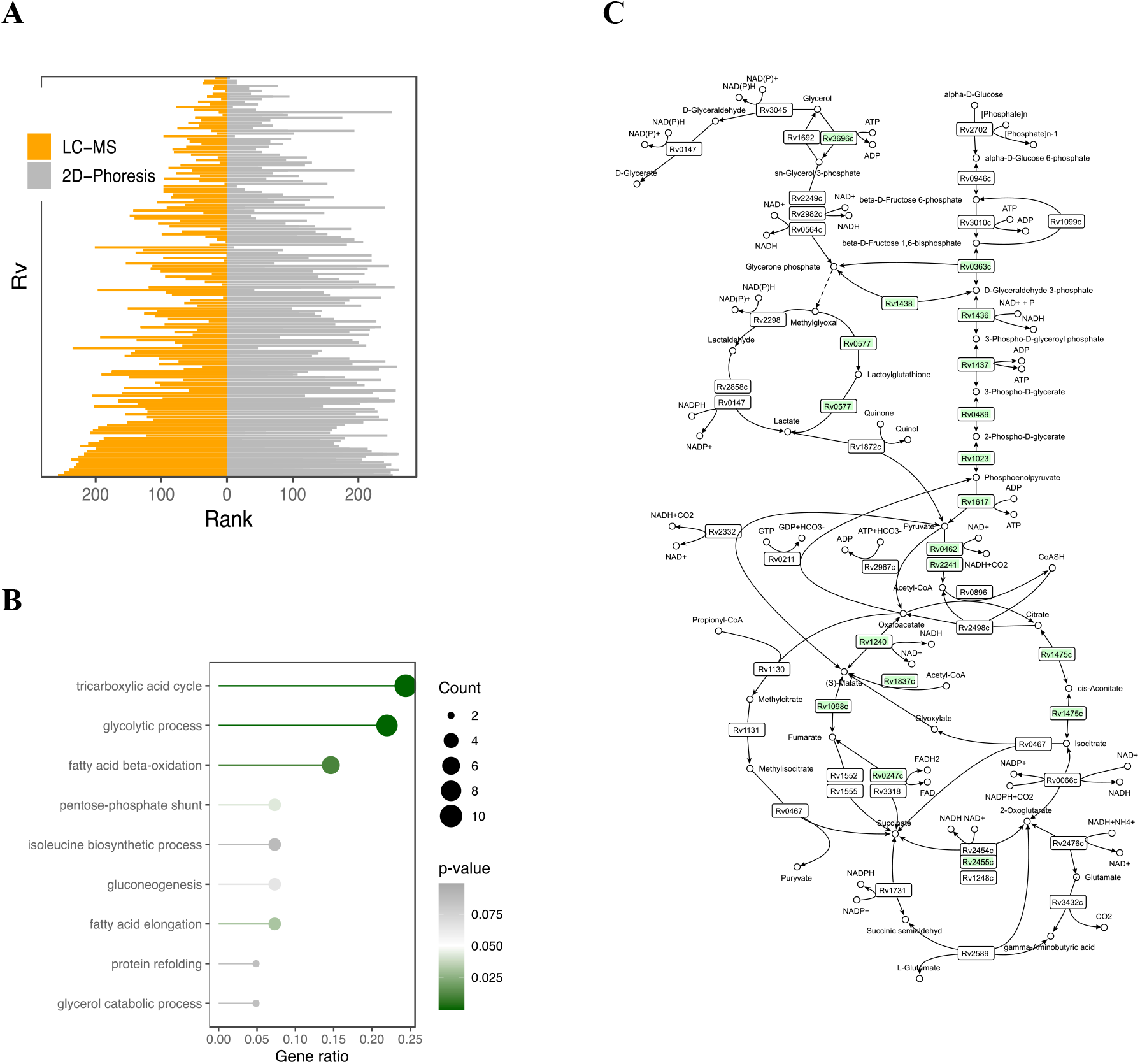
Comparison of the current LC-MS data with the previously published work on 2D proteomics of dormant *M. tuberculosis* (16). A) Visualization of the results of Mann–Whitney comparison of data acquired in the current study with the previously published data on 2D-proteomic analysis; B) GO enrichment analysis for the pathways assembled by the enzymes stored in D1; C) coverage of central carbon metabolism in D1.

The high-throughput LC-MS-based reconstruction of central carbon metabolism significantly increased the coverage of metabolic processes, and additionally disclosed mycothiol and a *de novo* UMP biosynthetic pathway (Fig. 5). The annotated pathways resemble the adjustment of mycobacterial metabolic machinery for life inside lipid-rich macrophages, particularly involvement of the methylcitrate and glyoxylate pathways to utilize the excess acetyl- and propionyl-CoA (resulting from the catabolic oxidation of fatty acids, cholesterol and branched amino-acid catabolism) (60–63). It is worth noting that such biochemical adaptation is tightly linked to mycobacterial virulence; thus, isocitrate lyase, besides its biochemical functions of participation in the glyoxylate cycle, in the methyl citrate cycle of propionyl-CoA metabolism or in succinate-associated generation for maintenance of proton-motive force in dormancy (14, 64), is an important mycobacterial virulence factor (60, 65). The decrease in virulence in *icl* mutants is precisely related to the enzymatic function of Icl and reflects the importance of the glyoxylate shunt for bacterial life *in vivo*.

The prevalence of catabolic enzymes in the absence of respiration may result in the generation of an excess of reduced FADH_2_ and NADH, which should be reoxidized for the continuation of metabolism (66). In the absence of respiration, a proposed mechanism for NAD^+^ replenishment and maintaining the energized level of transmembrane potential consists of exploiting the reductive branch of the TCA cycle (a part of the Arnon–Buchanan cycle) (47, 64), which is corroborated by the recently published data on succinate accumulation in the same model by dormant *M. smegmatis* cells (14).

Nonetheless, the current data on relative protein changes may provide only speculative information on real flux flows inside the cells (67), albeit speculatively envisaging metabolic adaptation of the cells to persistence alongside the activation of virulence factors, favouring the host’s immune escape. At the same time, enzymes stored in dormancy that are involved in particular metabolic pathways may be exploited by dormant cells during reactivation, when macromolecular biosynthetic processes are initiated (68). For example, the salvage of enzymes involved in the metabolism of purine and pyrimidine, the key precursors of DNA and RNA, in the current study (similarly to the observation of accumulation of the corresponding metabolites in the metabolism of dormant *M. smegmatis* (14)) makes these precursors immediately accessible after the onset of reactivation.

At the same time, individual enzymes preserved in dormancy can retain functionality, in particular those with cytoprotective functions.

Thus, under excess exogeneous glycerol one would expect the accumulation of carbonyl compounds, e.g., lactaldehyde (14) or methylglyoxal (Fig. 5). Therefore, it is to be expected that mycobacteria have developed mechanisms of protection from such reactive electrophilic agents.

Functioning NAD(P)H-dependent aldehyde dehydrogenases ensure the flow of metabolic transformations of carbonyl compounds and, on the other hand, may represent valuable targets for TB drug discovery (69). The glyoxylase Rv0577, presumably ensuring the detoxification of methylglyoxal, has similarly been considered as a target for pyrimidine imidazoles (70).

Whereas a majority of organisms (including human beings) exploit glutathione as a carbonyl detoxification system, mycobacteria lack glutathione, being dependent on mycothiol, a conjugate of N-acetylcysteine with *myo*-inosityl-2-amino-deoxy-α-D-glucopyranoside (71). Aside from its antioxidant properties, mycothiol may act as a cofactor for aldehyde dehydrogenase and participate in detoxification of rifampicin and isoniazid (71, 72). Correspondingly, pathway enrichment analysis disclosed the salvage of biosynthetic enzymes of mycothiol biosynthesis (Fig. 5E).

Part of the adaptation to oxidative stress is the accumulation of F_420_-dependent enzymes: glucose-6-phosphate dehydrogenase – fgd1 (Rv0407) and F_420_H_2_-dependent quinone reductase (deazaflavin-dependent nitroreductase, Rv1261c). The first passes electrons from the resulted in gluconeogenesis glucose-6-phosphate on F_420_H_2,_ which further participates in specific two-electron reduction of quinones by the F_420_H_2_-dependent quinone reductase (deazaflavin-dependent nitroreductase, Rv1261c), preventing the one-electron reduction pathway and the formation of cytotoxic semiquinones, notorious for formation of superoxide radical. The evolution of this self-protective mechanism was necessary, considering menaquinone as a main quinone of mycobacteria, which due to its more negative redox potential than ubiquinones tends to generate superoxide (73).

Thus, alongside the individually functioning SODs, catalases and chaperones, mycobacterial cells are armoured with efficient defence systems, protecting the bacteria against various stress factors.

Therefore, the current model of self-acidification of *Mtb* in the post-stationary phase being an *in vitro* model, it mirrors the virulence and biochemical adaptation of these bacteria to *in vivo* persistence. Comparative analysis of proteins the stored in dormancy found in the present study with the recently established immunogenic *Mtb* proteins resulted in 40 proteins with immunogenic properties (Fig. 6). Despite the serodiagnosis of latent TB being very obscure, the work of Zhou et al. (32), where serodiagnostic analysis was carried out using the sera of patients with confirmed latent TB at the proteome level, is of particular interest. Among 62 seropositive TB proteins reported by Zhou et al., 12 proteins out of 40 were found in the current study. Additionally, experiments recently carried out on the sera of 42 TB patients allowed the detection of a set of 27 potentially immunoreactive proteins (74), six of them (Rv2623, Rv2018, Rv2145c, Rv2744c, Rv1837c, Rv0341) similarly detected in the curent study with a positive FC in the dormant state. It is to be anticipated that the list of selected antigenic proteins includes those that may be of interest for identifying the specific ‘immunosignature’ of latent TB, which can later be used for immunodiagnosis of latent TB. However, this assumption requires further experimental verification.

The currently available therapy with the commonly exploited ‘first- and second-line’ TB drugs is directed at a rather narrow spectrum of *Mtb* targets, controlling the processes of DNA replication and transcription translation or targeting the enzymes of cell-wall biosynthesis/remodelling. However, all these ‘targets’ require the active flow of physiological pathways in the bacterial cells, sparing therefore the population of metabolically inactive dormant cells. Therefore, accounting for the urgent need for new chemicals efficient against drug-resistant mycobacteria and drug-tolerant dormant populations, it is an attractive idea to consider the enzymes available in dormancy capable of biotransformation, which could activate prodrugs and result in the accumulation of substances with general toxicity (ROS, RNS and chemically reactive metabolites and antimetabolites) (11, 12).

Some prodrugs such as clofazimine or delamanid which are activated by intracellular specific cofactor-dependent enzymes may serve as an illustration of this approach. The first molecule generates intracellular ROS through NADH dehydrogenase (54). The latter undergoes a metabolic conversion by mycobacterial deazaflavin-dependent nitroreductase followed by the release of intracellular NO, eradicating therefore a dormant *Mtb* subpopulation (75, 76).

Knowledge of the rare cofactors and cofactor-dependent oxidoreductases accumulated in dormancy may contribute to the development of new redox compounds (Supplement). Additionally, the bioactivation of prodrug compounds in hydrolytic processes, saved intact in dormancy (Supplement), is potent for the development of new anti-latent TB chemicals. Preliminary success of this concept has been demonstrated for nitazoxanide and methotrexate, albeit the particular bioactivating hydrolases were not disclosed (11). Malfunctioning of the reactive carbonyl detoxification systems under a surplus of glycerol may lead to the accumulation of (methyl)glyoxal/lactaldehyde in dormant mycobacteria (14). *In vivo*, a fat-rich diet may result in the accumulation of similar toxic carbonyl compounds within the cells: glyceraldehyde, glyceronphosphate, methylglyoxal, crotonaldehyde, malondialdehyde, 4-hydroxynonenal and, therefore, the application of inhibitors of the carbonyl detoxifying process may lead to inactivation of the bacteria (77, 78). On the other hand, lipid catabolism may elicit ‘ketone body’ intracellular acidosis (in the case of surplus acetyl-CoA and malfunction), in particular of malate synthase (Rv1837c) (79). A possible malate synthase inhibitor, mimicking the structure of methyl glyoxal – phenyl-diketo acid inhibitor – was recently proposed (80) and its applicability for the inactivation of dormant mycobacteria looks promising.

In conclusion, the current study demonstrated the biochemical adaptation of *Mtb* to the conditions of ‘non-culturability’ and visually demonstrated involvement of the pH-dependent regulator phoP in the suppression of components of the DOS regulon and in the activation of a number of processes crucial for cellular survival. Further, biochemical processes essential for the development and support of dormancy (methylcitrate and glyoxylate pathways, the salvage of mycothiol and UMP biosynthetic pathways as well as protective systems) were revealed. In the present study we, for the first time, annotate enzymes significantly abundant in dormant *Mtb* cells (oxidoreductases and hydrolases), which potentially may be further considered as biotransformatory enzymes – prodrug activators suitable for the elimination of dormant mycobacteria.

## Acknowledgements

This work was funded by the Russian Science Foundation – Grant 19-15-00324

## Author Contributions

VN carried out statistical and bioinformatics analysis: analysed and visualized data, prepared the figures, wrote the manuscript. MS – designed the project, supervised the project, provided the cells and cellular extracts for the analysis; DL, FD, JS – carried out proteomic profiling; annotated mycobacterial proteins. AK – supervised the project, corrected the draft.

The Authors read, commented and approved the final version of the manuscript.

